# FTO-mediated m6A demethylation of ULK1 mRNA promotes autophagy and activation of hepatic stellate cells in liver fibrosis

**DOI:** 10.1101/2024.03.14.584975

**Authors:** Tingjuan Huang, Chunhong Zhang, Junjie Ren, Qizhi Shuai, Xiaonan Li, Xuewei Li, Jun Xie, Jun Xu

**Affiliations:** Shanxi Key Laboratory of Birth Defect and Cell Regeneration, Department of Biochemistry and Molecular Biology, Shanxi Medical University, Taiyuan, 030001, Shanxi, China; Department of Gastroenterology and Hepatology, The First Hospital of Shanxi Medical University, Taiyuan, China; Department of Cancer Radiotherapy Department, Shanxi Provincial People’s Hospital, Taiyuan, 030001, China; Department of Hepatopancreatobiliary Surgery, The First Hospital of Shanxi Medical University, Taiyuan, 030001, China

**Keywords:** Autophagy, hepatic stellate cells, FTO, m^6^A methylation, ULK1

## Abstract

The activation of hepatic stellate cells (HSCs) is the central link in the occurrence and development of liver fibrosis. Our previous studies showed that autophagy promotes HSCs activation and ultimately accelerates liver fibrosis. Unc-51-like autophagy activating kinase 1 (ULK1) is an autophagic initiator in mammals and N^6^-methyladenosine (m^6^A) modification is closely related to autophagy. In this study, we find that m^6^A demethylase fat mass and obesity-associated protein (FTO) is upregulated during HSCs activation and bile duct ligation (BDL)-induced hepatic fibrosis, which is the m^6^A methylase with the most significant difference in expression. Importantly, we identify that FTO overexpression aggravates HSCs activation and hepatic fibrosis via autophagy. Mechanistically, compared with other autophagy-related genes, ULK1 is the target of FTO due to FTO mainly mediates the m^6^A demethylation of ULK1 and upregulates its expression, thereby enhancing autophagy and activation of HSCs. Noteworthy, m^6^A reader YTH domain-containing protein 2 (YTHDC2) decreases ULK1 mRNA level via recognizing the m^6^A binding site and ultimately inhibits autophagy and activation of HSCs. Taken together, our findings highlight m6A-dependent ULK1 as an essential regulator of HSCs autophagy and reveal ULK1 as a novel potential therapeutic target for hepatic fibrosis treatment.

**Graphical Abstract:** m6A demethylases FTO promoted autophagy via recognizing the ULK1 m6A binding site, thus triggering HSCs activation, and eventually leading to liver fibrosis. In this process, YTHDC2 participated in the translation of ULK1.

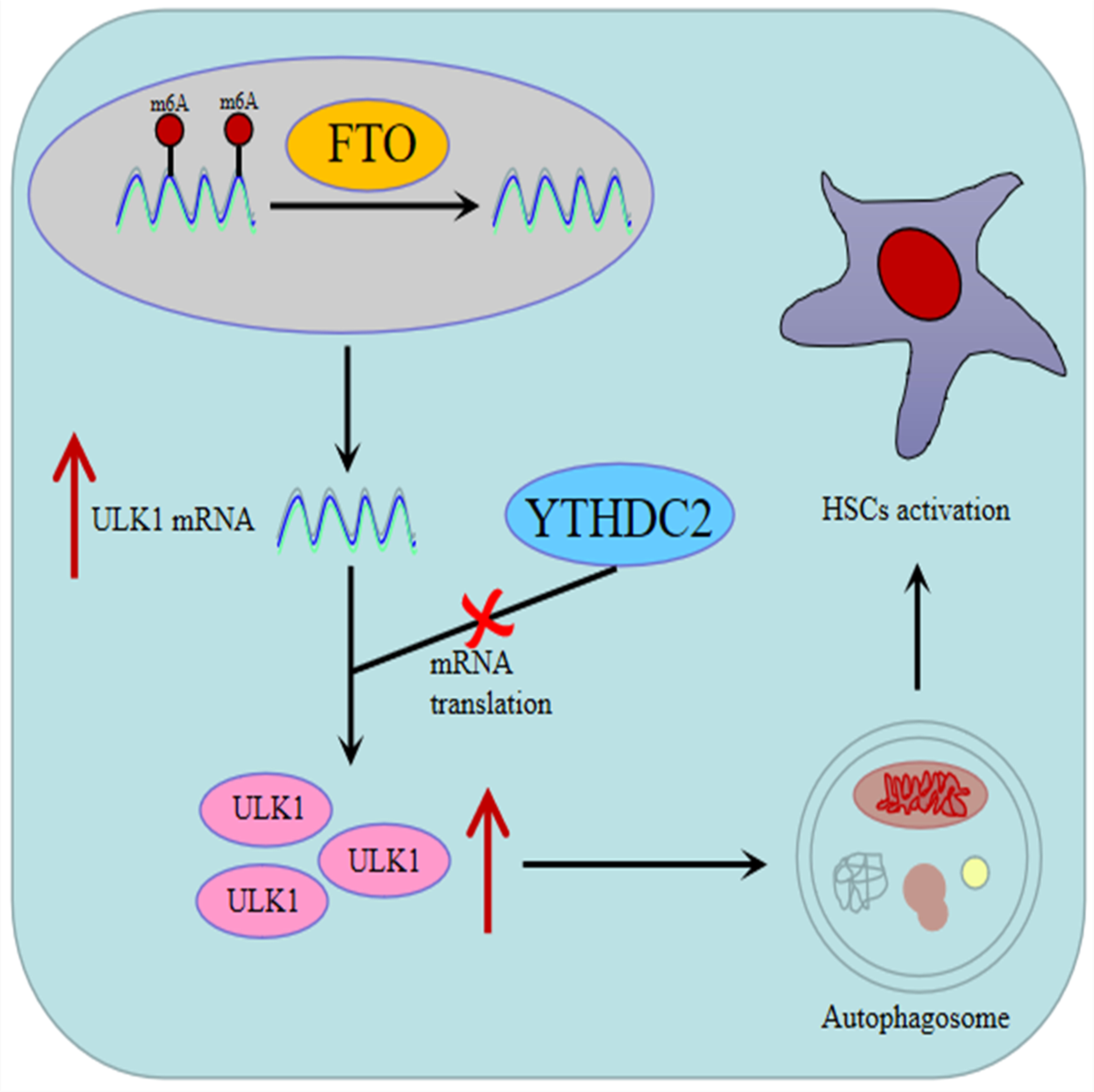

## 1. Introduction

Liver fibrosis is a complex physiological and pathophysiological condition following long-term chronic liver injury and is a necessary stage for various types of liver disease to develop into cirrhosis, liver failure and even liver cancer [1]. Cholestasis is an important factor of liver fibrosis, affecting approximately 200-500 individuals per million inhabitants [2]. Currently, in addition to improving the etiology and reducing cholestasis, there are still some limitations in the treatment of anti-hepatic fibrosis. The cholestatic liver disease is characterized by damage to the small intrahepatic biliary ducts, followed by proliferation and inflammatory response, which lead to the activation of hepatic stellate cells (HSCs). Hence, the elimination of activated HSCs is recognized to be the main strategy for anti-hepatic fibrosis.

Autophagy is a specific type of cell death, which starts with the formation of double-membrane autophagosomes, and ultimately fuses with the lysosomal compartments to degrade cellular organelles and proteins [3]. In hepatic fibrosis, autophagy plays a dual role. A report showed that impairing autophagy could promote nonalcoholic liver fibrosis progression [4]. Additional studies demonstrated that autophagy in HSCs attenuated CCL4-induced liver fibrosis [5]. We previously reported that autophagy was upregulated in bile duct ligation (BDL)-induced hepatic fibrosis, causing activation of HSCs, which ultimately accelerated liver fibrosis. Interestingly, a large amount of autophagy related genes (ATGs) are involved in the formation of autophagy, however, the specific ATGs regulating autophagy in cholestatic liver fibrosis remain unclear.

N^6^-methyladenosine (m^6^A) is the prominent dynamic mRNA modification, which is modulated by methyltransferase complex (writers), demethylases (erasers) and RNA-binding proteins (readers), influencing mRNAs stability, translation, subcellular localization, and alternative splicing [6]. The “writers” proteins containing WTAP (Wilms’ tumour 1 associated protein), METTL3 (methyltransferase like 3), METTL4, METTL14 catalyze m^6^A methylation. In contrast, ALKBH5 (alkB homolog 5) and FTO (obesity-associated protein) function as the “eraser” proteins to reverse m^6^A modifications. Additionally, m^6^A reader proteins, such as YTHDC1/2 (YTH-domain-containing protein 1/2) and YTHDF1/2/3 (YTH-domain-family 1/2/3), can bind to the m^6^A motif to affect RNA stability or function [7]. Importantly, exploring the post transcriptional regulation of m^6^A methylation on autophagy is helpful to develop new diagnostic and therapeutic targets for cholestatic hepatic fibrosis.

Autophagy genes are closely associated with m^6^A modification. A recent study reported that m^6^A modification controlled autophagy through upregulating ULK1 (Unc-51-like autophagy activating kinase 1) protein abundance in Hela cells [8], while another study showed that m^6^A methylation controlled autophagy and adipogenesis by targeting ATG5 and ATG7 in 3T3-L1 preadipocytes [9], indicating that the ATGs targeted by m^6^A methylation regulation may be distinct in different cells. However, it is unknown that which ATG is targeted by m^6^A methylation to induce autophagy and activation of HSCs and cholestatic hepatic fibrosis.

In the present study, we explored the potential mechanisms of autophagy regulated by m^6^A in HSCs activation. We found that m^6^A demethylases FTO promoted autophagy via recognizing the ULK1 m^6^A binding site, thus triggering HSCs activation, and eventually leading to liver fibrosis. Our study indicated that m^6^A modification may be a novel and important post-transcriptional regulator of HSCs autophagy and ULK1 may be a potential therapeutic target for hepatic fibrosis treatment.

## 2. Materials and methods

### 2.1 Animal experiments

The male C57BL/6 mice (20-22g) were maintained and bred in normal condition. The animal study was reviewed and approved by the Institutional Animal Care and Use Committee of Shanxi Medical University, and the license Key was SCXK2021-0006. Two liver fibrosis models were required. Briefly, common bile duct ligation (BDL) was performed during laparotomy under isoflurane inhalation anesthesia by isolating and ligating the common bile duct (n=8 per group). In sham animals, only surgery was performed without ligation of the common bile duct (n=8 per group). According to our previous reports[10], cholestatic liver fibrosis formed approximately 4 weeks after the operation, and liver tissues were harvested and stored in formalin and liquid nitrogen. Another liver fibrosis model was formed by transducing FTO (0.1 ml of each at a titer of 2*10^9^ pfu) into liver tissue via the tail vein with adenovirus. The recombinant adenoviral vector encoding FTO (Ad-FTO) was synthesized by Sangon Biotech (Shanghai). This group of mice were sacrificed at 12 weeks (n=8), and liver tissues were harvested.

### 2.2 Cell culture and treatment

Immortalized mouse HSC JS-1 (provide by the Experimental Center of Science and Research of the First Hospital of Shanxi Medical University) were cultured in DMEM with 10%FBS, 100U/ml penicillin, and 100ug/ml streptomycin at 37□ in an incubator with 5% CO_2_. When JS-1 cells were fused to about 80% in adherent culture, AdFTO was transfected for 24h, and then the cells were starved with serum-free DMEM for another 24h. After FTO siRNA (Sangon, Shanghai) was transfected into JS-1 cells with LipoFiter^TM^ 3.0 (Hanbio, Hunan) for 6h, the cells were cultured under starvation until 48h. To investigate whether FTO regulates autophagy and activation of HSCs by targeting ULK1, the cells were added Ad FTO for24h after the start of ULK1 siRNA transfection for 6h, followed by starvation treatment for 24h. Each group in vitro was performed in triplicate.

### 2.3 Antibodies

Primary antibodies against α-SMA (ab124964, 1:1000), collagen I (ab260043, 1:1000), LC3B (ab48394, 1:1000), SQSTM1/P62 (ab91526, 1:1000) were obtained from Abcam Technology. FTO (sc-271713 1:500,) was obtained from Santa Cruz.

### 2.4 Histological analyses

Liver tissue samples were paraffin embedded and 4 μm tissue sections stained with standard methods and subjected to histopathological analysis according to our reports previously [10]. Briefly, with 4% paraformaldehyde to fix, xylene to dewax, alcohol to hydrate, and boiled sodium citrate buffer to repair antigen. Hepatic morphology and liver fibrosis were examined by H&E, Masson and Immunohistochemistry staining. For Immunohistochemistry staining, the sections were incubated with primary antibodies at 4 □ overnight and with secondary antibodies at 37 □ for 20 min. Finally, staining was detected using DAB (Beyotime).

### 2.5 Western blot

BCA colorimetric method was used to quantify the protein concentration. 4 x SDS loading buffer (P1015, Solarbio) were added in samples and boiled for 10 min. Equal amounts of protein samples were fractionated by electrophoresis through agarose gels and transferred to PVDF membranes (Bio-Rad). After blocking with 5% dry milk for 2h, the membranes were incubated with primary antibodies at 4 □ overnight and secondary antibodies at room temperature for 1-2 h. All the primary antibodies were diluted at a concentration of 1:1000. Proteins were detected by enhanced chemiluminescence (ECL; Bio-Rad).

### 2.6 RNA extraction and quantitative real-time PCR

RNA was extracted using Trizol solution (15596018, Invitrogen) per the manufacturer’s instructions, and one μg of RNA was reverse transcribed using a cDNA Reverse Transcription Kit (RR037A, Takara). Final cDNA samples were used then used for quantitative real-time PCR assay by SYBR Green PCR Kit (RR820A, Takara). Quantitative primers were designed based on the gene sequences by Sangon Biotech. The target gene primer sequences are provided in Table 1.

**Table 1.**
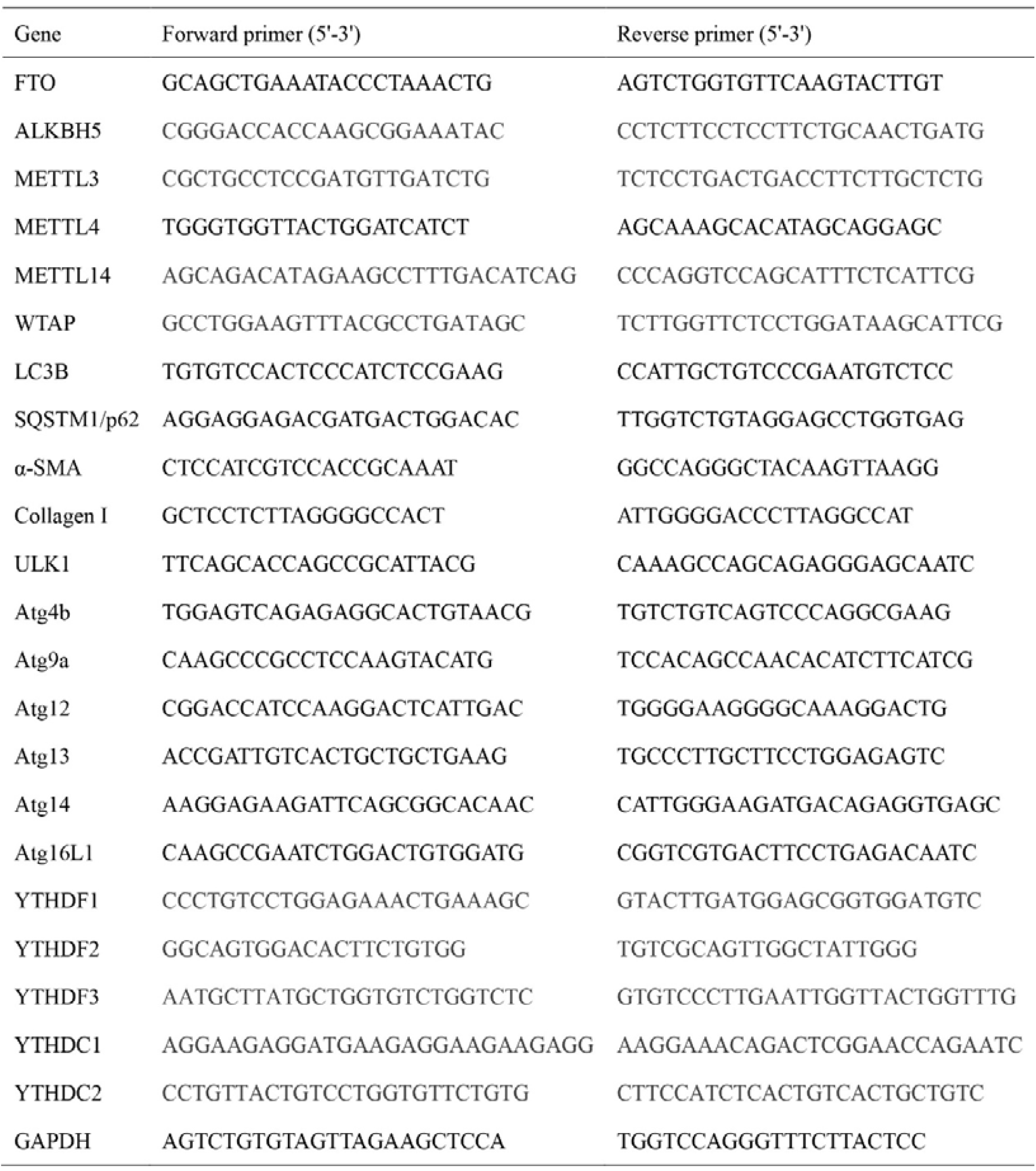
Primers sequences for RT-qPCR.

### 2.7 Immunofluorescence

The cells were fixed with 4% paraformaldehyde for 15-20 mins, permeabilized with 0.1% Triton X 100 for 15 mins, and then blocked with 1% BSA at room temperature for 30 mins. Next, cells were incubated with primary antibodies at 4 □ overnight followed by incubation with FITC labeled secondary antibodies at room temperature for 1 h. DAPI (C0065, Solarbio) was used to stain nuclei. An inverted fluorescence microscope was then used to observe its fluorescence.

### 2.8 MeRIP qPCR

Magna MeRIP^TM^ m^6^A Kit (17-10499, Merck) was used to enable identification and transcriptome-wide profiling of m^6^A. Firstly, RNA is chemically fragmented into 100 nucleotides or smaller fragments followed by magnetic immunoprecipitation with a monoclonal antibody toward m^6^A. After immunoprecipitation, isolated RNA fragments can be subjected with RT-qPCR.

### 2.9 m^6^A-seq

The m6A-specific antibody is most often necessary in high-throughput sequencing approaches for m6A-sequencing. The m6A sequencing was performed in Guangzhou Kidio Biotechnology. Total RNA were extracted with Trizol, fragmented and immunoprecipitated by m6A antibody according to Magna MeRIP^TM^ m^6^A Kit, and then isolated RNA fragments can be subjected with RNA sequencing.

### 2.10 Quantification of m^6^A RNA

For detecting m^6^A RNA methylation status directly, total RNA was isolated and were quantified using EpiQuik m^6^A RNA Methylation Quantitative kit (P-9005-48, Epigentek). Relative m^6^A RNA methylation levels were then calculated using the formula provided by the manufacturer’s instructions.

### 2.11 Transmission electron microscopy

The cells were fixed with 2% glutaraldehyde for 2 h. After washing with sodium arsenate for several hours, the cells was fixed with 1% osmic acid for more than 2 h. Using a gradient of acetone for dehydration at 5-15 min per stage, and the mixture of acetone and embedding agent was soaked at room temperature for 2-4 h. The epoxy resin was embedded for 24 h at 40□ and polymerized for 48h at 60□. Ultrathin slices were obtained with an ultramicrotome (LKB), placed in uranium acetate and lead citrate dye for 15-30 min, transmission electron microscopy was performed.

### 2.12 Statistical analysis

All data were expressed as mean ± standard error of the mean (SEM), and all calculations were performed using GraphPad Prism 8.0 or SPSS 23.0 statistical software. In addition, Statistical significance was evaluated using Student’s t-test or one-way analysis of variance. P-value<0.05 was considered statistically significant.

## 3. Results

### 3.1 FTO is upregulated during HSCs activation and BDL-induced hepatic fibrosis

Similar to our previous research methods that cholestatic liver fibrosis was induced by BDL [11]. HE of livers from BDL treated mice showed increase in dilation of the bile duct, proliferation of small bile ducts, as well as infiltration of inflammatory cells in the portal area and around the bile duct, and Masson staining showed collagen deposition in BDL group (Fig. 1A). Moreover, immunohistochemical staining showed that BDL increased expression of α-SMA (Fig. 1B). These results implied the BDL-induced hepatic fibrosis model was successfully established. In addition, to explore the potential possibility of m^6^A modification in cholestatic liver fibrosis, we first employed qPCR analysis to compare the mRNA expression of m^6^A writers and erasers. Interestingly, the results showed that FTO were the most obvious increase in BDL-induced hepatic fibrotic tissue with upregulated protein and mRNA (Fig. 1C, D).

**Figure 1.**
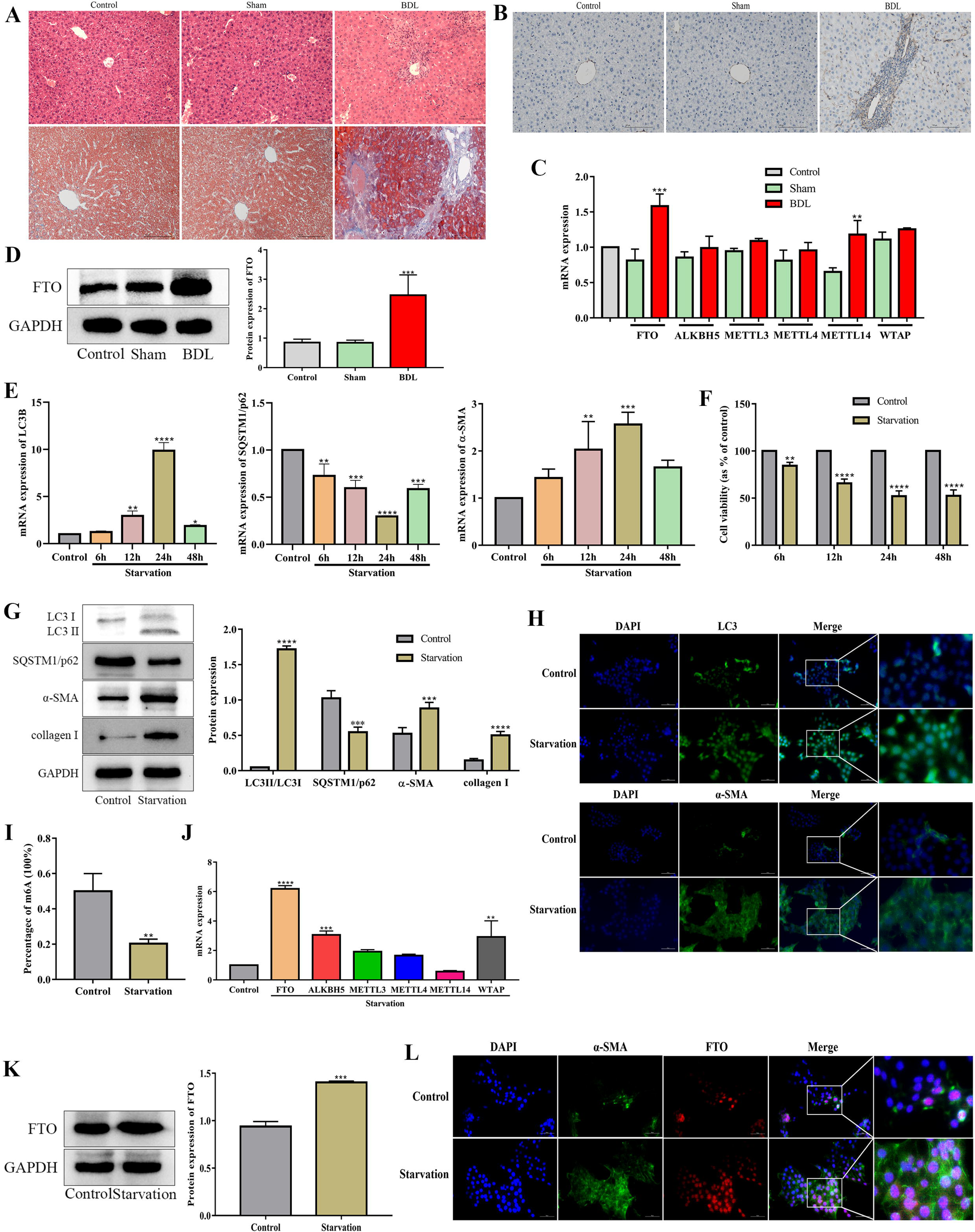
FTO is upregulated during HSCs activation and BDL-induced hepatic fibrosis. (A) HE and Masson staining were performed to evaluate the establishment of BDL-induced hepatic fibrosis model. Scale bars: 100µm. (B) α-SMA was semiquantitative detected by immunohistochemical staining. Scale bars: 100µm (**P<0.01 vs. Sham). (C) mRNA level of m6A writers and erasers were detected by RT-qPCR in BDL-induced hepatic fibrosis (**P<0.01 vs. Sham). (D) Protein expression of FTO with western blot in BDL-induced hepatic fibrosis (**P<0.01 vs. Sham). (E) mRNA expression of autophagy and HSCs activation with RT-qPCR in JS-1 cells with starvation treatment at different time points (**P<0.01 vs. Control). (F) Effect of starvation on viability of JS-1 cells (**P<0.01 vs. Control). (G) Protein levels of autophagy and HSCs activation with western blot in JS-1 cells with starvation treatment (**P<0.01 vs. Control). (H) Immunofluorescence of LC3B and α-SMA in JS-1 cells with starvation treatment. The cells were stained for LC3B (green), α-SMA (green) and DAPI (blue). Scale bars: 20µm. (I) The m6A levels were detected by m6A RNA Methylation Quantitative kit (**P<0.01 vs. Control). (J) mRNA level of m6A writers and erasers were detected by RT-qPCR in vitro (**P<0.01 vs. Control). (K) Protein levels of FTO with western blot in JS-1 cells with starvation treatment (**P<0.01 vs. Control). (L) Co positioning of FTO and α-SMA: Immunofluorescence of FTO and α-SMA in JS-1 cells with starvation treatment. The cells were stained for FTO (red), α-SMA (green) and DAPI (blue). Scale bars: 20µm. All data are presented as the mean ± SD (n=8 in vivo, n =3 in vitro).

HSCs activation is the central link in the occurrence and development of liver fibrosis. In order to confirm whether FTO is involved in HSCs activation, we verified it in vitro. Autophagy is known to occur during starvation [12, 13]. Consistent with literature reports [14], our results showed that autophagy was increased by Starvation and peaked at 24h in JS-1 cells (Fig. 1E) while cell vitality was decreased (Fig. 1F), which was used for subsequent experiments. Western blot showed the ratio of LC3 II/I was upregulated, while p62 was downregulated in starvation group at 24h, further suggesting that autophagy was enhanced by starvation (Fig. 1G). Meanwhile, the expression of α-SMA and collagen I were dramatically increased by starvation, indicating starvation induced HSCs activation (Fig. 1G). Immunofluorescence showed the same results (Fig. 1H). Subsequently, we examined the levels of m^6^A modification during HSCs activation, m^6^A RNA methylation quantification assay showed a significant decline in m^6^A levels after treatment with starvation (Fig. 1I). As expected, consistent with the results in vivo, FTO remained the most obvious increase in Hunger-induced HSCs activation (Fig. 1J, K). Finally, we conducted immunofluorescence colocalization of FTO and α-SMA, the results showed that the main upregulated regions of FTO are highly coincident with HSCs regions (Fig. 1L). These results indicated that FTO may play an essential role in the process of HSCs activation.

### 3.2 FTO promotes HSCs activation and hepatic fibrosis as well as autophagy

To verify the role of FTO in hepatic fibrosis in vivo, we first transfected AdFTO into mouse liver. HE and Masson staining showed FTO enhanced inflammatory cell infiltration and collagen deposition in the portal area (Fig. 2A). Western blot showed that α-SMA and collagen I in AdFTO group were gradually upregulated (Fig. 2C), which was consistent with immunohistochemical staining (Fig. 2B). Our previous studies showed that enhanced autophagy level accelerated hepatic fibrosis[10]. In this study, we hypothesized that the autophagy was essential in FTO-induced hepatic fibrosis. To test the hypothesis, we detected the level of autophagy after FTO treatment. As expected, the results showed that the conversion of LC3 I to LC3 II in AdFTO group was gradually increased while p62 was decreased (Fig. 2C), suggesting that autophagy was enhanced by FTO.

**Figure 2.**
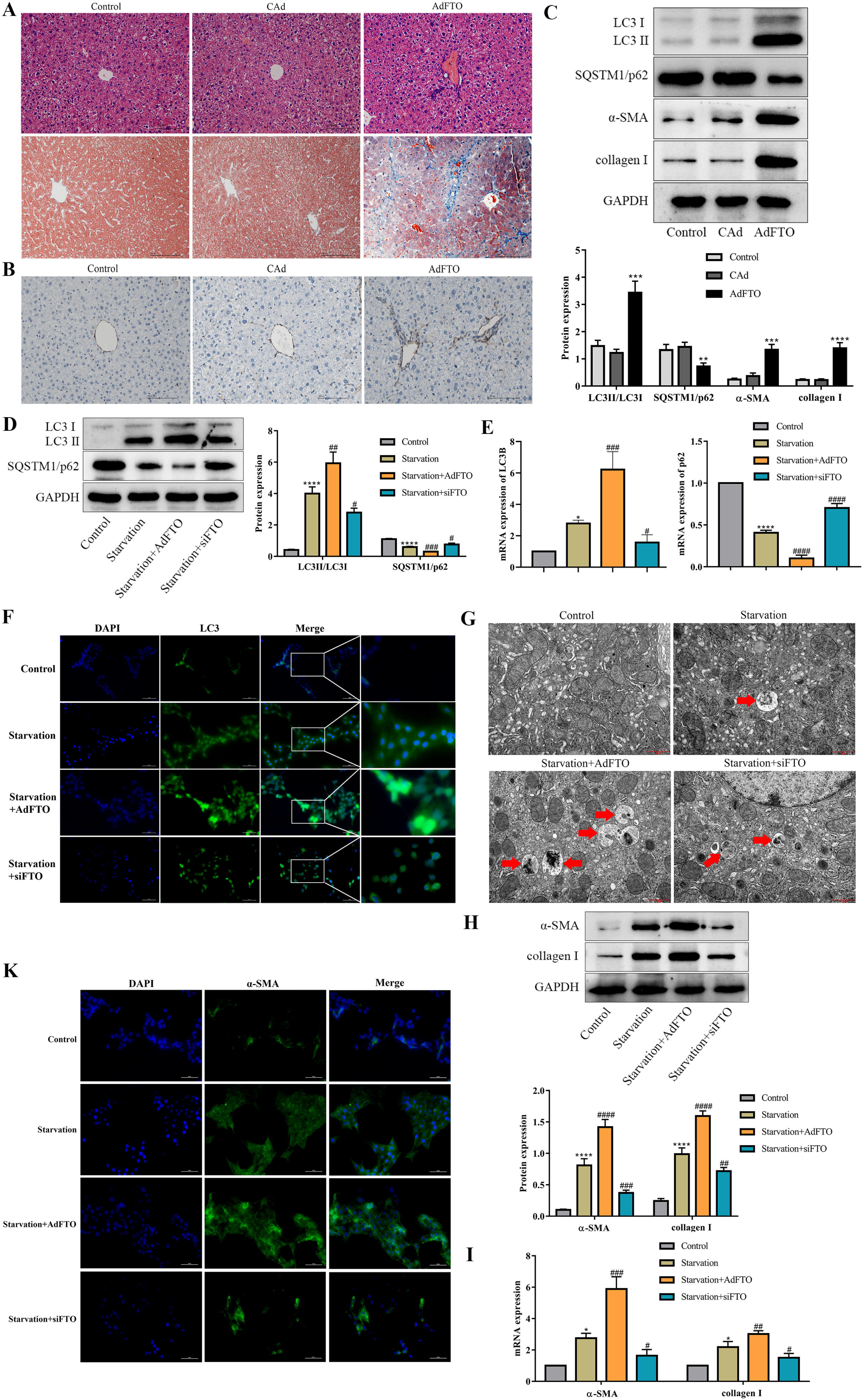
FTO promotes HSCs activation and hepatic fibrosis as well as autophagy. (A) The pathological changes of liver treated by FTO were monitored by HE and Masson staining. Scale bars: 100µm. (B) α-SMA of liver by FTO was semiquantitative detected by immunohistochemical staining. Scale bars: 100µm (**P<0.01 vs. CAd). (C) Protein expression LC3B, p62, α-SMA, collagen I with western blot in mouse livers transfected by AdFTO (**P<0.01 vs. CAd). (D, E) Protein and mRNA levels of autophagy with western blot and RT-qPCR in starvation induced JS-1 cells transfected by AdFTO or siFTO (**P<0.01 vs. Control, ^##^P<0.01 vs. Starvation). (F, K) Immunofluorescence of LC3B and α-SMA in starvation induced JS-1 cells transfected by AdFTO or siFTO. The cells were stained for LC3B (green), α-SMA (green) and DAPI (blue). Scale bars: 20µm. (G) Transmission electron microscopy was used to assess autophagosome. Scale bars: 1µm. (H, I) Protein and mRNA levels of α-SMA and collagen I with western blot and RT-qPCR in starvation induced JS-1 cells transfected by AdFTO or siFTO (**P<0.01 vs. Control, ^##^P<0.01 vs. Starvation). All data are presented as the mean ± SD (n=8 in vivo, n =3 in vitro).

Whether autophagy was involved in FTO-induced HSCs activation in vitro? To clarify this potential possibility, AdFTO and siFTO were transfected respectively on starvation-treated JS-1 cells. As shown in Fig. 2D, FTO markedly increased starvation-induced the ratio of LC3 II/LC3 I and decreased p62 expression while siFTO reversed these changes, which were consistent with RT-qPCR results (Fig. 2E). Meanwhile, immunofluorescence suggested that the level of LC3 was increased by AdFTO and decreased by siFTO (Fig. 2F). In addition, transmission electron microscopy confirmed that AdFTO increased the number of autophagic lysosomes while siFTO decreased that (Fig. 2G). Subsequently, the HSCs activation markers were further detected. The results suggested that the promotion effect of starvation on α-SMA and collagen I was further strengthened by AdFTO and weakened by siFTO (Fig. 2H, I). Immunofluorescence of α-SMA showed the same result (Fig. 2K). All these findings supported that FTO promoted HSCs autophagy and activation.

### 3.3 FTO promotes HSCs activation via autophagy

To verify whether autophagy was required for FTO-induced HSCs activation, autophagy inhibitor 3-MA was used in JS-1 cells in the presence of FTO. RT-qPCR showed that 3-MA significantly reduced FTO-induced LC3 upregulation and further increased FTO-induced p62 expression (Fig. 3A, B), indicating that 3-MA suppressed the autophagy inducing effect of FTO. To further understand how autophagy impacts HSCs activation, we examined the role of the autophagy inhibitor 3-MA in FTO-induced HSCs activation. The results showed that 3-MA treatment markedly reduced α-SMA and collagen I mRNA expression (Fig. 3C, D), suggesting that reducing autophagy affected the promoting effect of FTO on HSCs activation. Coimmunofluorescent staining of α-SMA and LC3 displayed the same result (Fig. 3E). Based on above results, autophagy was required to mediate FTO-induced HSCs activation.

**Figure 3.**
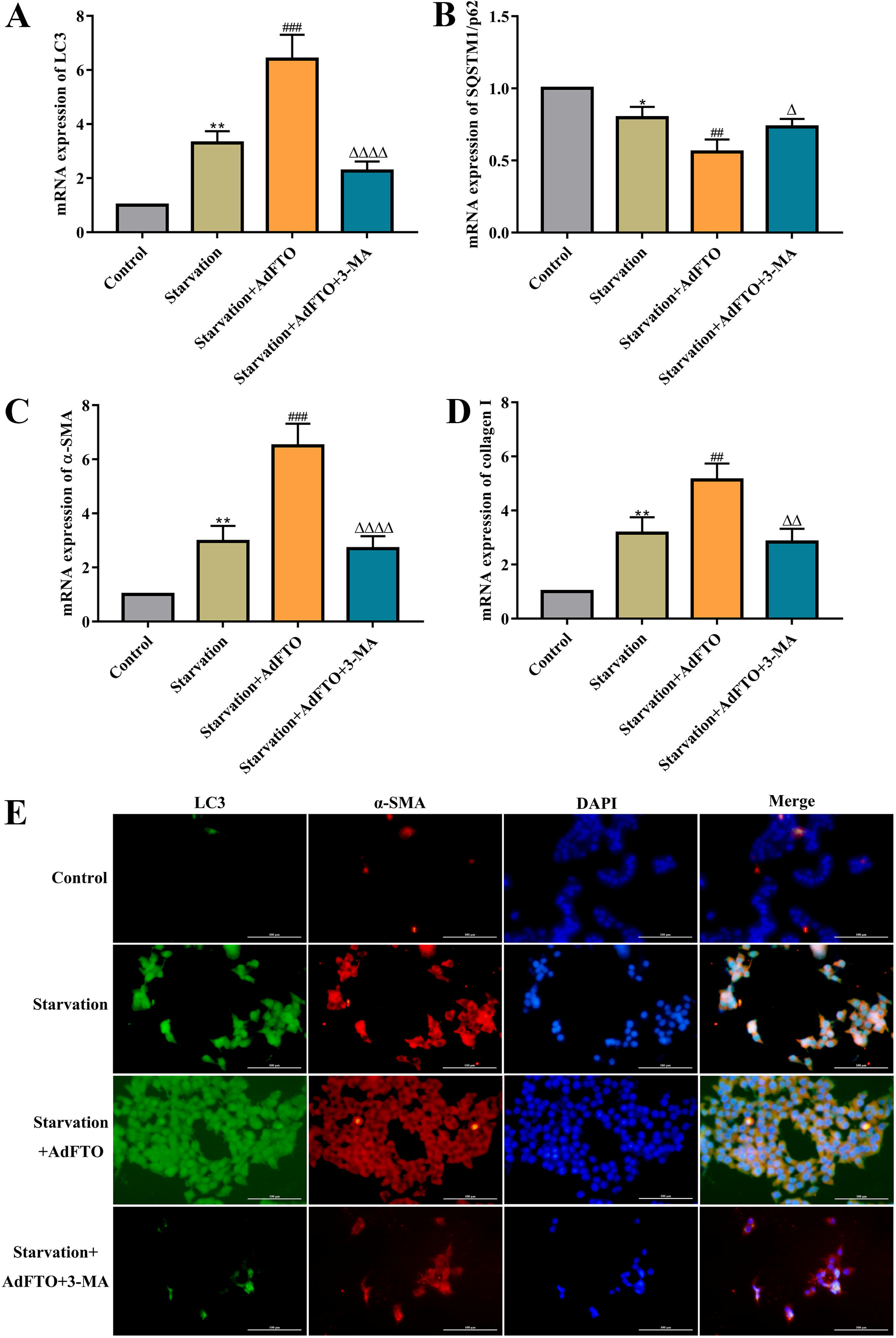
FTO promotes HSCs activation via autophagy. (A, B, C, D) mRNA levels of LC3, SQSTM1/p62, α-SMA and collagen I with RT-qPCR in starvation induced JS-1 cells after AdFTO treatment in the presence or absence of 3-MA. (E) Coimmunofluorescent staining of α-SMA and LC3 in starvation induced JS-1 cells after AdFTO treatment in the presence or absence of 3-MA. (**P<0.01 vs. Control, ^##^P<0.01 vs. Starvation, ^ΔΔ^P<0.01 vs. Starvation+AdFTO). All data are presented as the mean ± SD (n =3 in vitro).

### 3.4 FTO mainly mediates the m^6^A demethylation of ULK1 to drive autophagy

We performed m^6^A sequencing (m^6^A-seq) of major ATGs to further explore which ATGs have m^6^A modification in BDL-induced hepatic fibrosis. The results showed that a total of 8 ATGs have m^6^A modification sites, including ULK1, p62, ATG4b, ATG9a, ATG12, ATG13, ATG14, ATG16L1 (Fig. 4A). Then, to identify the target genes of FTO in autophagy, siFTO was transfected into JS-1 cells and all ATGs with m^6^A modification were detected by MeRIP-qPCR. The results indicated that knockdown of FTO increased the m^6^A level on mRNA transcripts of ULK1, p62, ATG9a, ATG13, ATG16L1, especially ULK1 significantly (Fig. 4B). More importantly, to deeply assess which ATG was the direct target of FTO, AdFTO and siFTO were transfected into JS-1 cells followed by starvation. Interestingly, we found that compared with starvation group, overexpression of FTO markedly upregulated protein levels of ULK1 while knockdown of FTO downregulated that (Fig. 4C). The other ATGs, such as ATG9a, ATG13, ATG16L1 were not significantly changed (Fig. 4C). The expression of p62 was opposite to that of ULK1 (Fig. 3C), further indicating that FTO mainly mediates the m^6^A demethylation of ULK1 to promote autophagy in starvation-induced HSCs activation.

**Figure 4.**
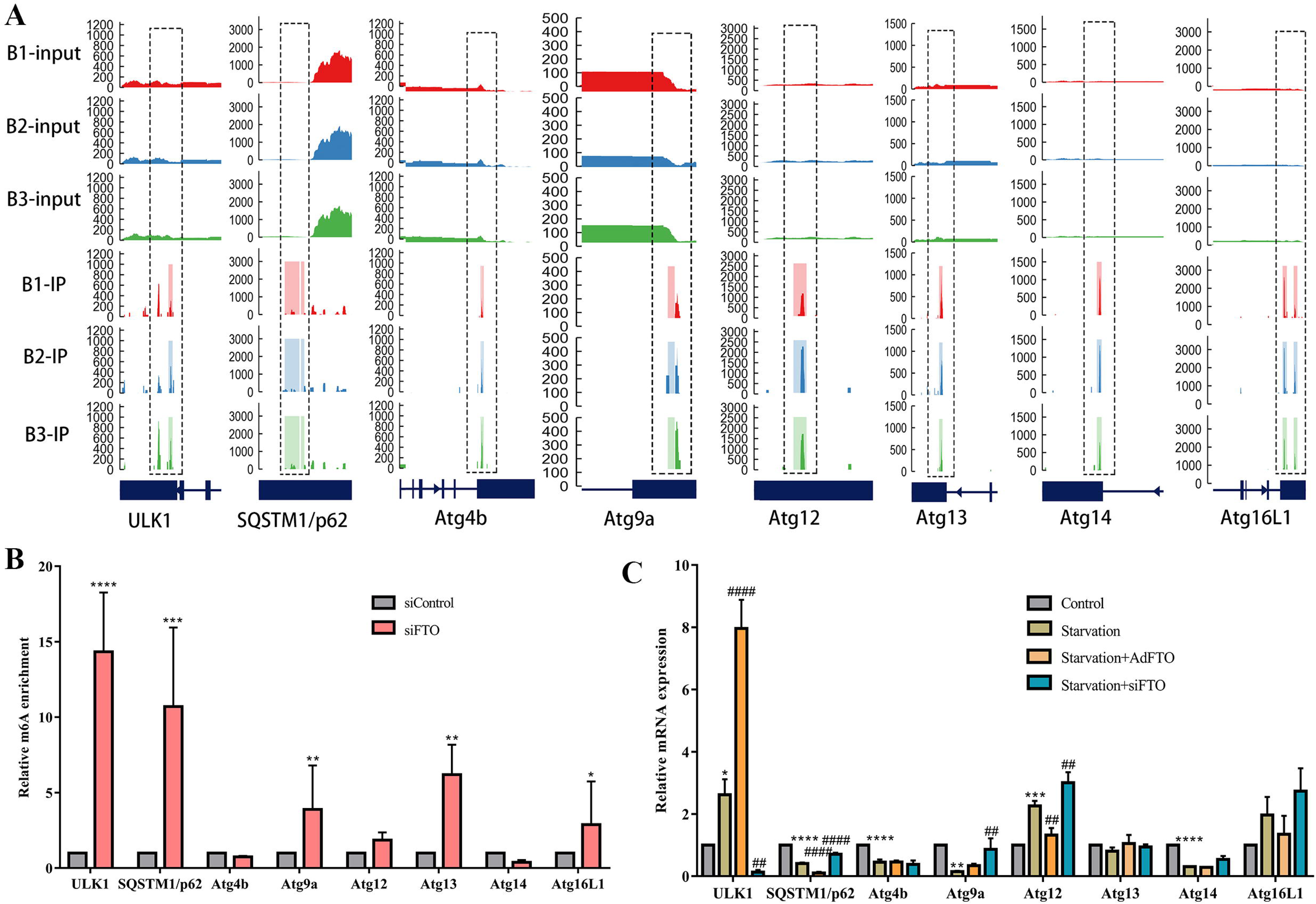
FTO mainly mediates the m6A demethylation of ULK1 to induce autophagy in HSCs activation. (A) m6A-seq of m6A peaks at ULK1, SQSTM1/p62, ATG4b, ATG9a, ATG12, ATG13, ATG14, ATG16L1 mRNAs. In the X-axis, blue boxes represent exons, and blue lines represent introns. The Y-axis shows sequence read number. (B) MeRIP-qPCR analysis of m6A levels of ULK1, SQSTM1/p62, ATG4b, ATG9a, ATG12, ATG13, ATG14, ATG16L1 mRNA in Control and siFTO cells (**P<0.01 vs. siControl). (C) mRNA expression of ULK1, SQSTM1/p62, ATG4b, ATG9a, ATG12, ATG13, ATG14, ATG16L1 in starvation induced JS-1 cells transfected by AdFTO and siFTO (**<0.01 vs. Control, ^##^P<0.01 vs. Starvation). Data are presented as the mean ± SD (n =3 in vitro).

### 3.5 FTO enhances autophagy and activation of HSCs by targeting ULK1

To confirm whether FTO promotes autophagy and HSCs activation by affecting the expression of ULK1, we performed rescue experiment and observed that silencing of ULK1 reversed the upregulation of LC3 II/LC3 I ratio and downregulation of the protein expression of p62 in FTO-overexpressing JS-1 cells followed by starvation (Fig. 5A). Similar to western blot results, RT-qPCR also showed the same outcome (Fig. 5B). In consistent, silencing ULK1 significantly reduced LC3 puncta augmented by FTO in immunofluorescence assays (Fig. 5C). Moreover, compared with starvation+AdFTO group, ULK1 siRNA decreased the number of autophagosomes (Fig. 5D). These results demonstrated that FTO regulated autophagy by targeting ULK1.

**Figure 5.**
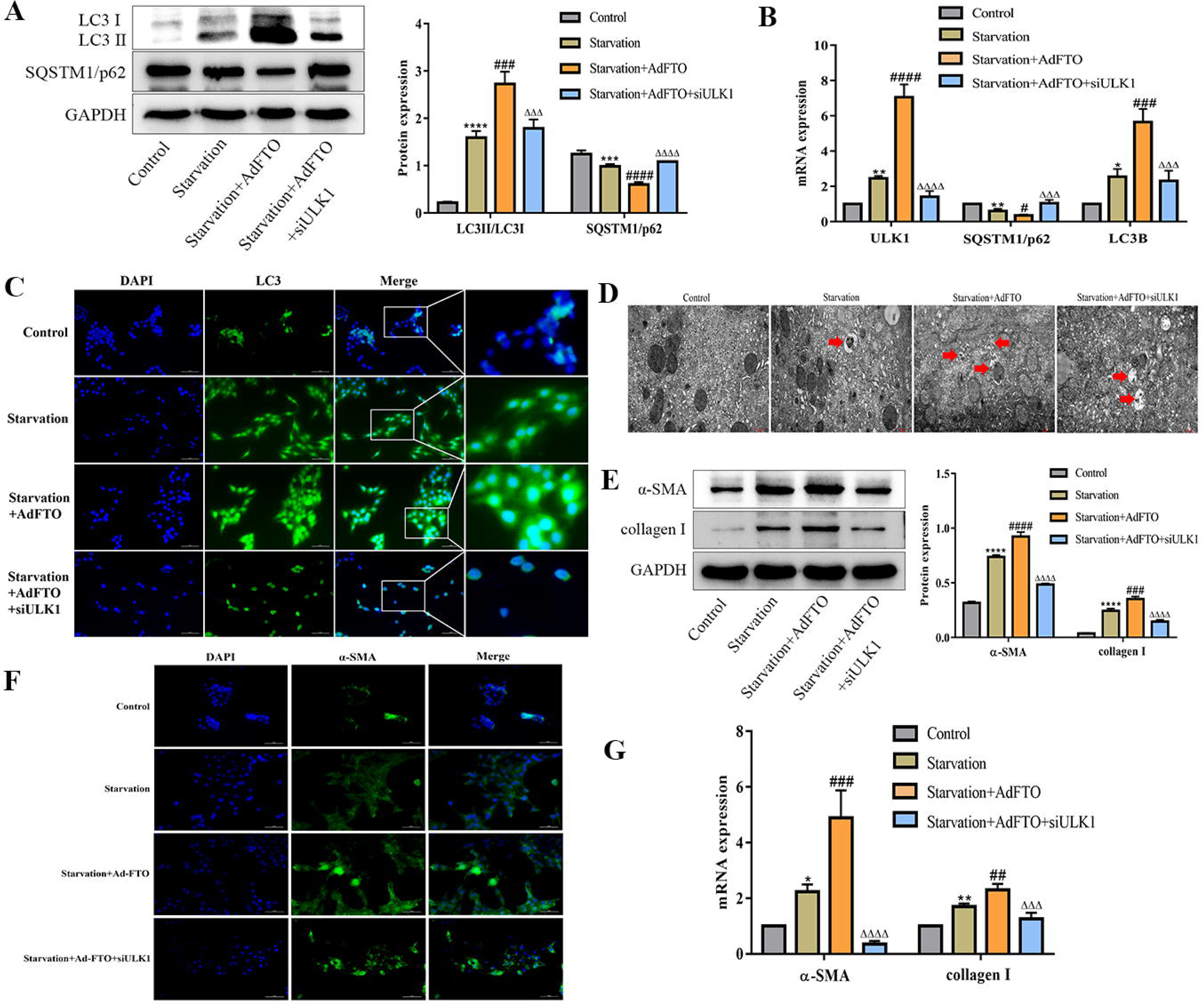
FTO promotes autophagy and activation of HSCs by targeting ULK1. (A, B) JS-1 cells treated with ULK1 siRNA were transfected with AdFTO for 24h, and then were treated with Starvation. The protein and mRNA levels of autophagy were measured by western blot and RT-qPCR (**P<0.01 vs. Control, ^##^P<0.01 vs. Starvation, ^ΔΔ^P<0.01 vs. Starvation+AdFTO). (C, F) JS-1 cells treated with ULK1 siRNA were transfected with AdFTO for 24h, and then were treated with Starvation. Immunofluorescence was used to detect LC3B and α-SMA. The cells were stained for LC3B (green), α-SMA(green) and DAPI (blue). Scale bars: 20µm. (D) Transmission electron microscopy was used to assess autophagosome. Scale bars: 1µm. (E, G) JS-1 cells treated with ULK1 siRNA were transfected with AdFTO for 24h, and then were treated with Starvation. The protein and mRNA of α-SMA and collagen I were measured by western blot and RT-qPCR (**P<0.01 vs. Control, ^##^P<0.01 vs. Starvation, ^ΔΔ^P<0.01 vs. Starvation+AdFTO). All data are presented as the mean ± SD (n =3 in vitro).

Next, we investigated whether ULK1 was involved in FTO-induced HSCs activation. As expected, the protein expression of α-SMA and collagen I increased by FTO were decreased upon ULK1 knockdown (Fig. 5E). In addition, knockdown of ULK1 decreased α-SMA puncta enhanced by FTO in immunofluorescence assays (Fig. 5F). Consistently, the mRNA levels of α-SMA and collagen I upregulated by FTO were downregulated upon ULK1 knockdown (Fig. 5G). Taken together, these results illustrated that FTO targeted ULK1 and mediated its expression in an m^6^A-dependent manner, and further regulated HSCs autophagy and activation.

### 3.6 YTHDC2 regulates ULK1 mRNA translation via recognizing the m^6^A binding site and further affects HSCs autophagy and activation

The YTH domain-containing proteins, including YTHDF1-3 and YTHDC1-2 participate in numerous RNA processes, such as mRNA splicing, nuclear export, translation and decay in post-transcriptional regulation. To further explain which YTH domain-containing protein mediates the regulation of autophagy by m^6^A modification, we preliminarily explored the expression of YTHDF1-3 and YTHDC1-2 in HSCs activation and in BDL-induced hepatic fibrosis. The results showed that YTHDC2 mRNA rather than YTHDF1-3 and YTHDC1 increased most significantly both in vitro and in vivo (Fig. 6A, B). It has been found that YTH domain-containing proteins play important roles in post-transcriptional modification process hence modulate the expression of target genes including autophagy [15]. In order to verify whether YTHDC2 bound to ULK1 mRNA in HSCs JS-1 cells, RNA IP was performed. As shown as in Fig. 6C, compared with GAPDH PCR product, the ULK1 PCR products were highly enriched in YTHDC2, indicating a direct bind of YTHDC2 to ULK1 mRNA. Furthermore, MeRIP-qPCR showed that the mRNA transcripts of ULK1 was significantly enriched by YTHDC2 (Fig. 6D). In addition, actinomycin D assay showed that YTHDC2 reduced the stability of ULK1 mRNA (Fig. 6E). Importantly, the expression of ULK1 mRNA and protein were significantly decreased by YTHDC2 (Fig. 6F, I), which is related to the translation function of YTHDC2 regulating ULK1.

**Figure 6.**
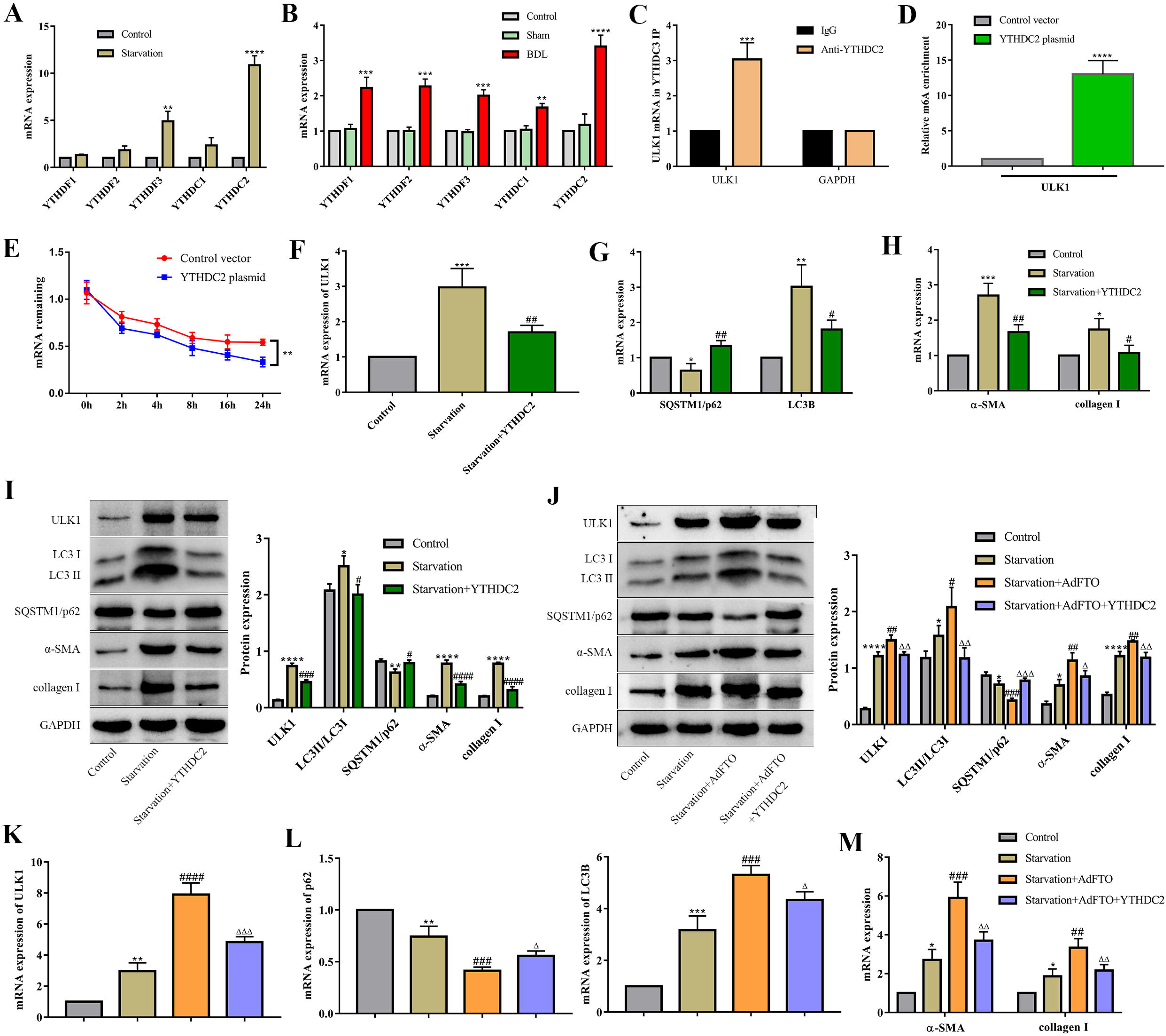
YTHDC2 regulates the expression of ULK1 and HSCs activation in an m6A-dependent manner. (A, B) mRNA level of m6A readers were detected by RT-qPCR in vitro and in vivo (**P<0.01 vs. Sham). (C) The binding of YTHDC2 and ULK1 mRNA was measured by ribonucleoprotein immunoprecipitation (RNA IP) (**P<0.01 vs. IgG group). (D) MeRIP-qPCR analysis of m6A levels of ULK1 mRNA in Control and YTHDC2 plasmid (**P<0.01 vs. Control). (E) RNA stability of ULK1 in Control and YTHDC2 plasmid (**P<0.01 vs. Control). (F, G, H, I) mRNA and protein expression of ULK1, LC3B, SQSTM1/p62, α-SMA and collagen I in starvation induced JS-1 cells transfected by YTHDC2 plasmid (**P<0.01 vs. Control, ^##^P<0.01 vs. Starvation). (J, K, L, M) JS-1 cells treated with YTHDC2 plasmid were transfected with AdFTO for 24h, and then were treated with Starvation. The protein and mRNA levels of ULK1, LC3B, SQSTM1/p62, α-SMA and collagen I were measured by Western blot and RT-qPCR (**P<0.01 vs. Control, ^##^P<0.01 vs. Starvation, ^ΔΔ^P<0.01 vs. Starvation + AdFTO). Data are presented as the mean ± SD (n =3 in vitro).

To confirm the effect of YTHDC2 on HSCs autophagy and activation, YTHDC2 plasmid was transfected into JS-1 cells followed by starvation. The results revealed that YTHDC2 downregulated the mRNA and protein expression of LC3 and upregulated p62 expression (Fig. 6G, I). As expected, α-SMA and collagen I were markedly decreased by YTHDC2 (Fig. 6H, I). To further proved that m^6^A of ULK1 recognized by YTHDC2 was related to FTO, HSCs was transfected with YTHDC2 plasmid followed by FTO. The results showed that YTHDC2 attenuated the upregulation of ULK1 induced by FTO (Fig. 6J, K). Similarly, the autophagy level and HSCs activation increased by FTO were also reversed by YTHDC2 (Fig. 6J, L, M). Collectively, these data demonstrated that YTHDC2 reduced ULK1 mRNA expression via recognizing the m^6^A binding site and further decreased HSCs autophagy and activation.

## 4. Discussion

The hepatic fibrosis model was induced through BDL. HE, Masson staining and high expression of α-SMA showed that the liver fibrosis model was successful. Accumulating evidences have shown that m^6^A-related enzymes regulate liver fibrosis, such as Li et al. revealed that Mettl3 deficiency attenuated HSCs activation and CCl_4_-induced liver fibrosis [16], and Sun et al reported that the m^6^A reader YTHDF3 alleviated liver fibrosis by CCl_4_ [17]. Furthermore, Li et al indicated that exenatide significantly reversed high-fat-induced lipid accumulation and inflammatory changes accompanied by decreased FTO expression[18]. In this study, we revealed d that compared with other methylases, FTO was upregulated during HSCs activation and BDL-induced hepatic fibrosis, which is the m^6^A methylase with the most significant difference in expression. In addition, the upregulation region of FTO was highly coincident with HSCs regions, indicating the increase of FTO is responsible for HSCs activation and cholestatic hepatic fibrosis.

The activation of HSCs is the central link in the occurrence and development of liver fibrosis. m^6^A modification has been reported to play a critical role in HSCs activation[19]. Attractively, Shen et al recently reported that overexpression of FTO evidently damaged dihydroartemisinin (DHA)-induced ferroptosis thus promoting HSCs activation [20]. Importantly, our study firstly identified that FTO overexpression aggravated HSCs activation and hepatic fibrosis, whereas FTO knockdown markedly inhibited it. Furthermore, Previous studies presented that autophagy was involved in the activation of HSCs in cholestatic liver fibrosis[11]. To clarify the underlying regulatory mechanisms of FTO on HSCs activation and liver fibrosis, autophagy was detected. Shen et al showed that overexpression of FTO reduced DHA-induced autophagy[20], However, in this study, we demonstrated that FTO upregulated the levels of autophagy in HSCs and ultimately increased the expression of activation markers in HSCs through autophagy, which was consistent with our previous study that aggravating autophagy promoted HSCs activation[10]. Therefore, downregulation of FTO mediated the upregulation of m^6^A modification may become a valid method to inhibit HSCs activation.

m^6^A is the most prevalent posttranscriptional modification of mRNA, occurring at approximately three to five sites per mRNA in mammals[21]. Accumulating studies have indicated that m^6^A modification participates the autophagy activation by modifying ATGs and further regulates a variety of physiological and pathological processes[22]. Interestingly, Ming et al demonstrated that Mettl14 regulates osteoclast differentiation via inducing autophagy through m^6^A /IGF2BPs/Beclin1 signal axis[23]. Moreover, He et al. revealed that METTL3 inhibits the apoptosis and autophagy of chondrocytes in inflammation through mediating m^6^A /YTHDF1/Bcl2 signal axis[24]. In the current study, we found that ULK1 was rich in methylation sites due to the increased m^6^A modification by demethylase FTO knockdown. Further, overexpression of FTO significantly upregulated the level of ULK1, while there were no significant difference in other ATGs. Our study suggested for the first time that FTO mainly mediated the m^6^A demethylation of ULK1 to drive autophagy.

Serving as a protein kinase, ULK1 is activated upon autophagy stimulation and is critical for recruiting other autophagy-related proteins to the autophagosome formation sites[25]. Liu et al verified that ULK1 activator BL-918 displays a therapeutic potential on amyotrophic lateral sclerosis through inducing cytoprotective autophagy[26]. Based on accumulating evidence, we further confirmed the role of ULK1 in FTO induced autophagy and activation of HSCs. Indeed, our data suggested that ULK1 was positively correlated with HSCs autophagy and activation. More importantly, ULK1 knockdown reduced the formation of autophagosome and ECM accumulation, which were markedly increased by FTO. There results provide evidence that FTO enhances autophagy and activation of HSCs by targeting ULK1.

Recently, the m^6^A reader proteins were found to participate in post-transcriptional modification process thus regulating the expression of related genes involved in inflammatory and autophagy[15]. According to a report, m^6^A reader protein YTHDF1 regulated autophagy by targeting SQSTM1 in diabetic skin[27]. In our study, it was found that YTHDC2 was the most significant differential gene among YTH family in HSCs activation and cholestatic liver fibrosis. Then we confirmed that overexpression of YTHDC2 decreased ULK1 level via recognizing the m^6^A binding site and ultimately inhibited autophagy and activation of HSCs. More importantly, YTHDC2 weakened the increase of ULK1 induced by FTO, thereby improving autophagy and HSCs activation raised by FTO. However, the mechanism how YTHDC2 modulates ULK1 thus affecting HSCs activation needs to be further explored.

## 5. Conclusions

In summary, the study demonstrated that m^6^A demethylase FTO promotes HSCs autophagy by targeting ULK1, resulting in HSCs activation, and ultimately leading to hepatic fibrosis. Further, the m^6^A reader YTHDC2 is identified to specifically regulates ULK1. In brief, our findings provide a new insight into that targeting FTO/ULK1 axis may be a promoting strategy for clinical anti-liver fibrosis therapy.

## Acknowledgements

This study was supported by Fundamental Research Program of Shanxi Province (20210302124615), Research Fund for the Doctor Program of Shanxi Province (SD1913), Research Fund for the Doctor Program of Shanxi Medical University (XD1913) and National Nature Science Foundation of China (82203221).

## Author contributions

T-J H designed, performed the research and wrote the manuscript. C-H Z and J-J R performed the research and analyzed the data. Q-Z S and X-N L collected the literature. X-W L annotated and maintained the research data. J X and J X supervised the research and revised the manuscript. All authors read and approved the final manuscript.

## Data Availability Statement

The authors confirm that the data supporting the findings of this study are available within the article.

## Ethical approval

The animal study was reviewed and approved by the Institutional Animal Care and Use Committee of Shanxi Medical University, and the license Key was SCXK2021-0006.

## References

[1] Zhang J, Li Y, Liu Q, et al. Sirt6 Alleviated Liver Fibrosis by Deacetylating Conserved Lysine 54 on Smad2 in Hepatic Stellate Cells. Hepatology. 2021;73(3):1140–1157.

[2] Gossard AA, Talwalkar JA, Cholestatic liver disease. Med Clin North Am.2014;98(1):73–85.

[3] Klionsky D, Autophagy: from phenomenology to molecular understanding in less than a decade. Nature reviews. Molecular cell biology. 2007;8(11):931–7.

[4] Baselli G, Jamialahmadi O, Pelusi S, et al. Rare ATG7 genetic variants predispose patients to severe fatty liver disease. 2022;77:596–606.

[5] Gao J, Wei B, de Assuncao T, et al. Hepatic stellate cell autophagy inhibits extracellular vesicle release to attenuate liver fibrosis. 2020;73:1144–54.

[6] Fu Y, Dominissini D, Rechavi G, et al. Gene expression regulation mediated through reversible mLJA RNA methylation. Nature reviews. Genetics. 2014;15(5):293–306.

[7] Wu R, Jiang D, Wang Y, et al. N (6)-Methyladenosine (m(6)A) Methylation in mRNA with A Dynamic and Reversible Epigenetic Modification. Mol Biotechnol.2016;58(7):450–9.

[8] Zhao X, Yang Y, Sun BF, et al. FTO-dependent demethylation of N6-methyladenosine regulates mRNA splicing and is required for adipogenesis. Cell Res. 2014;24(12):1403–19.

[9] Deng X, Su R, Weng H, et al. RNA N(6)-methyladenosine modification in cancers: current status and perspectives. Cell Res.2018;28(5):507–517.

[10] Jin S, Zhang X, Miao Y, et al, m(6)A RNA modification controls autophagy through upregulating ULK1 protein abundance. Cell Res.2018;28(9):955–957.

[11] Wang X, Wu R, Liu Y, et al. m(6)A mRNA methylation controls autophagy and adipogenesis by targeting Atg5 and Atg7. Autophagy.2020;16(7):1221–1235.

[12] Huang TJ, Ren JJ, Zhang QQ, et al, IGFBPrP1 accelerates autophagy and activation of hepatic stellate cells via mutual regulation between H19 and PI3K/AKT/mTOR pathway. Biomed Pharmacother. 2019;116:109034.

[13] Kong YY, Huang TJ, Zhang HY, et al. The lncRNA NEAT1/miR-29b/Atg9a axis regulates IGFBPrP1-induced autophagy and activation of mouse hepatic stellate cells. Life Sci. 2019;237:116902.

[14] Duckney P, Li C, Hussey P, et al. TraB, a Novel Plant ER-Mitochondrial Contact Site Protein Functions as a Mitophagy Receptor in Plants. Autophagy. 2022;29;1–3.

[15] González-Rodríguez P, Delorme-Axford E, Bernard A, et al. SETD2 transcriptional control of ATG14L/S isoforms regulates autophagosome-lysosome fusion. Cell Death Dis. 2022;13(11):953.

[16] Chai W, Ye F, Zeng L, et al. HMGB1-mediated autophagy regulates sodium/iodide symporter protein degradation in thyroid cancer cells. Journal of experimental & clinical cancer research: CR. 2019;38(1):325.

[17] Liu S, Li G, Li Q, et al. The roles and mechanisms of YTH domain-containing proteins in cancer development and progression. American journal of cancer research. 2020;10(4):1068–1084.

[18] Li Y, Kang X, Zhou Z, et al. The m6A methyltransferase Mettl3 deficiency attenuates hepatic stellate cell activation and liver fibrosis. Molecular therapy. 2022;30(12):3714–3728.

[19] Sun R, Tian X, Li Y, et al. The m6A reader YTHDF3-mediated PRDX3 translation alleviates liver fibrosis. Redox biology. 2022;54:102378.

[20] Li S, Wang X, Zhang J, et al. Exenatide ameliorates hepatic steatosis and attenuates fat mass and FTO gene expression through PI3K signaling pathway in nonalcoholic fatty liver disease. Brazilian journal of medical and biological research. 2018;51(8):e7299.

[21] Yang J, Wang J, Yang Y, et al. ALKBH5 ameliorated liver fibrosis and suppressed HSCs activation via triggering PTCH1 activation in an mA dependent manner. European journal of pharmacology. 2022; 922:174900.

[22] Shen M, Li Y, Wang Y, et al. N-methyladenosine modification regulates ferroptosis through autophagy signaling pathway in hepatic stellate cells. Redox biology. 2021;47:102151.

[23] Shen M, Guo M, Li Y, et al. m(6)A methylation is required for dihydroartemisinin to alleviate liver fibrosis by inducing ferroptosis in hepatic stellate cells, Free Radic Biol Med. 2022;182:246–259.

[24] Lee M, Kim B, Kim V, et al. Emerging roles of RNA modification: m(6)A and U-tail. Cell. 2014;158(5):980–987.

[25] Liu L, Wang J, Sun G, et al. mA mRNA methylation regulates CTNNB1 to promote the proliferation of hepatoblastoma. Molecular cancer. 2019;18(1):188.

[26] He M, Lei H, He X, et al. METTL14 Regulates Osteogenesis of Bone Marrow Mesenchymal Stem Cells via Inducing Autophagy Through m6A/IGF2BPs/Beclin-1 Signal Axis. Stem Cells Transl Med. 2022;11(9):987–1001.

[27] He Y, Wang W, Xu X, et al. Mettl3 inhibits the apoptosis and autophagy of chondrocytes in inflammation through mediating Bcl2 stability via Ythdf1-mediated mA modification. Bone. 2022;154:116182.

